# Identification of short peptides that correlate with cytoplasmic retention of human proteins

**DOI:** 10.1101/2025.11.03.686273

**Authors:** Jay C. Brown, Baomin Wang

## Abstract

One group of human proteins found in the cytoplasm, but not in the nucleus is characterized by the presence of short (6-9aa), specific amino acid sequences thought to be involved in retaining proteins in the cytoplasm (cytoplasmic retention sequences). While strong evidence supports the ability of some peptides to act in this way, the number of such supported cases is small. We have taken the view that the situation would be improved by enhancing the methods available to identify cytoplasmic retention (CR) peptides. Here we describe an appropriate bioinformatic method to identify CR peptides using information about their location at the ends of cytoplasmic proteins. The method was then used to link seven different human cytoplasmic proteins with peptides suggested to have cytoplasmic retention activity. Further analysis was carried out with isoforms of the cytoplasmic proteins identified. Amino acid sequence information showed that while the proposed CR amino acid sequences can be the same or distinct in different protein isoforms, they are always located at the same site in the protein. For instance, while the proposed retention sequence of CCDC57 isoform X18 is MLARLVSNS, in isoform 7 it is SEPALNEL yet the two sequences are each located between amino acids 5 and 13 in the CCDC57 sequence. The results support the view that protein isoform is involved in determining the location of the CR sequence in a protein while the peptide sequence itself affects other variables such as the sub-region of the cytoplasm the protein needs to occupy. Overall, the study yielded identification of 15 candidate CR peptides in which 10 of the 15 have un-related amino acid sequences.

## 1. Introduction

For cell biologists it is of the utmost importance to understand factors that influence the subcellular location of proteins. A newly-synthesized protein must first be trafficked to the nucleus or the cytoplasm and thereafter to the appropriate sub-region such as the endoplasmic reticulum or mitochondria. Highly specific signals must be involved at each step, and if a protein needs to change its location because of developmental or environmental events, then additional signals and interpretation of signals must be involved. Much of the cell’s machinery must be devoted to delivering its proteins to the right places, and investigators have made it a high priority to clarify the factors involved.

A landmark in the field occurred in the early 1980s when highly specific amino acid sequences were found to be able to direct proteins to the nucleus [1-2]. Identification of similar features that influence localization to the cytoplasm, however, are complicated by the fact that proteins are synthesized in the cytoplasm making their transport to the cytoplasm unnecessary. Nevertheless, important advances have been made in identifying features that prevent proteins from leaving the cytoplasm, the cytoplasmic retention signals. For instance, a four amino acid sequence at the C-terminus of grp78 (HSPA5) has been found to mediate endoplasmic reticulum retention localization in COS cells [3]. Studies with fusion proteins involving a reporter protein linked to a candidate cytoplasmic retention sequence have been a rich source of information about the identity of cytoplasmic retention sequences [4-7]. Ankyrin repeats in NF-κB have been demonstrated to have cytoplasmic retention function [8].

We have taken the view that it would be an advance if investigators had access to a rapid method to identify cytoplasmic retention protein sequences. This could be used as a first step in identifying protein-protein contacts that underlie cytoplasmic structure and function. Here we describe a bioinformatic method designed to identify amino acid sequences able to confer cytoplasmic retention (CR) function on their home protein. Short amino acid sequences proposed for analysis were first tested for their presence in human proteins using NCBI-BLAST. Positive proteins were then screened further for the presence of the peptide at the C- or N-terminus and for protein location in the cytoplasm. Peptides with all three properties are suggested to have CR activity.

The above method was then used to test the amino acid sequence MLPRLVLNS for CR activity in human proteins and three candidate proteins were identified. The same analysis was performed with a second candidate peptide and with isoforms of all positive proteins. The experimental pathway resulted in the identification of fifteen novel candidate CR sequences.

## 2. Materials and Methods

### Short peptides chosen for analysis

Initial studies were carried out with two short peptides, MLPRLVLNS and MLARLVSNS. The two were selected because preliminary analysis demonstrated their relative abundance, compared with other short peptides, among human proteins.

### Blast analysis

NCBI BLAST (https://blast.ncbi.nlm.nih.gov/Blast.cgi) was used to identify homologous sequences in test peptides and human proteins. Options employed were blastp and ClusteredNR. Sequence identities were considered to be those where six or more contiguous sites are each occupied by the same amino acid. Analysis was pursued only with proteins where the homologous region is at or near the C- or the N-terminus because these sites are expected to be exposed on the protein surface and able to interact with other components of the cytoplasm.

### Other methods

The cellular location of individual human proteins were those given by Gene Cards (https://www.genecards.org/), Human Protein Atlas (https://www.proteinatlas.org/) and DeepLoc2.1 (https://services.healthtech.dtu.dk/services/DeepLoc-2.1/). Peptide structures were predicated by AlphaFold Server (https://alphafoldserver.com/welcome/). Peptide sequences were randomized with Sequence Manipulation Suite (https://www.bioinformatics.org/sms2/).

## 3. Results

### Study beginning with MLPRLVLNS

The overall project consisted of two parallel studies, each beginning with one of the two main test peptides, MLPRLVLNS and MLARLVSNS. In each case BLAST was used to probe the human genome for amino acid sequences matching the test peptide. Proteins with matches of six contiguous amino acids or more were then further examined for the presence of the other two features expected of a CR sequence, (1) location of the candidate peptide at or near the C- or N-terminus of the matching protein and (2) location of the protein in the cytoplasm. Proteins with all three properties are suggested to have cytoplasmic retention function.

Table 1 shows the results of the study carried out beginning with MLPRLVLNS. Proteins with sequences matching MLPRLVLNS or regions of it are shown in the “protein” column lines 1-18. Three proteins, MAP3K5, TRPC1 and TTPA (lines 1-3), were found to have the other two properties described above and are therefore proposed as proteins with CR activity. While proteins in lines 4-18 have matches with the test sequence, all fail one or both of the other CR properties and are not included in the CR candidate group. BCLC7, for instance, has a good match with the test protein sequence, but the matching sequence is not found at an end of the protein and the protein is found in the nucleus, not the cytoplasm (see line 4). While MYCBPAP has a good matching sequence located near a protein end (PRLVLNS; see line 15), the matching sequence is also found in CNOT6L, a protein present in the nucleus (line 16). Neither protein is therefore included with the list of CR candidates.

**Table 1:** Identification of human proteins with proposed cytoplasmic retention sequence MLPRLVLNS.

|  |  |  |  |  |  |  |
| --- | --- | --- | --- | --- | --- | --- |
| line | peptide sequence | protein | isoform | protein length (aa) | peptide location in protein | protein location in cell |
| 1 | MLPRLVLNS | MAP3K5 | X9 | 777 | 751-759 | cytosol, ER |
| 2 | MLPRLVLNS | TRPC1 | 16 | 308 | 17-25 | PM, cytosol, ER |
| 3 | MLPRLVLNS | TTPA | 4 | 282 | 274-282 | cytosol |
| 4 | LPRLVLNS | BCL7C | X1 | 324 | 183-190 | nucleus |
| 5 | RLVLNS | SLC28A1 | 4 | 543 | 536-541 | PM, cytosol |
| 6 | RLVLNS | KLRC3 | E | 240 | 235-240 | PM, cytosol |
| 7 | MLPRLV | DUOXA1 | 1 | 483 | 434-439 | ER, PM |
| 8 | MLPRLV | TTC2 | 2 | 372 | 346-351 | cytosol, nucleus |
| 9 | RLVLNS | TRMT10B | X5 | 285 | 262-267 | cytosol, nucleus |
| 10 | MLPRLV | GVQW3 | d | 187 | 161-166 | cytosol, nucleus |
| 11 | MLPRLV | TERF2IP | CRA_b | 161 | 1-6 | nucleus, cytosol |
| 12 | MLPRLV | NEU3 | e | 139 | 108-113 | lysosome |
| 13 | MLPRLV | ZNHIT1 | CRA_b | 129 | 1-6 | nucleus, cytosol |
| 14 | MLPRLV | ETFA | 2 | 45 | 19-24 | mitochondria, golgi |
| 15 | PRLVLNS | MYCBPAP | CRA_a | 855 | 33-39 | endosome, PM |
| 16 | PRLVLNS | CNOT6L | 5 | 587 | 299-305 | cytosol, nucleus |
| 17 | RLVLNS | ALG8 | 8 | 557 | 191-196 | ER, PM |
| 18 | MLPRLV | EIF4E | 2 | 248 | 139-144 | cytosol, nucleus |
|  | Randomize sequence: |  |  |  |  |  |
| 19 | LMLLPNVRS |  |  |  |  |  |
| 20 | LLPNVR | ADAMYS9 | 4 | 1907 | 639-644 | ER, PM |
| 21 | LPNVRS | CWF19L2 | none | 894 | 700-705 | nucleus |
| 22 | LPNVRS | GK | d | 559 | 230-235 | cytosol, nucleus |
|  | Rubinfeld et al. [6] |  |  |  |  |  |
| 23 | YLEQYYDPS | MAPK3 | 1 | 379 | 329-336 | golgi, cytosol |

ER: endoplasmic reticulum; PM: plasma membrane

A control experiment was performed with a randomized version of the test peptide (see line 19). BLAST analysis was performed with the randomized sequence in the expectation that it would yield fewer matches with human proteins and fewer of the other properties expected of candidate CR peptides. The expected result would help validate use of the test peptide to identify novel CR-active sequences. The outcome supported the expected resuilt (Table 1, lines 20-22). While 18 human proteins had matches with the authentic test sequence, only 3 were observed with the randomized version. Also, none of the matches with the randomized sequence were found at an end of its home protein.

Support for the bioinformatic approach used here comes from a study in which CR activity was found to be conferred by a short amino acid sequence of the human protein ERK2 (MAPK3) [6]. The active peptide was found near the C-terminus of the target gene as required of candidate CR peptides in the analysis reported here (Table 1, line 23).

### Study beginning with MLARLVSNS

Results of the study beginning with MLARLVSNS are shown in Table 2. They show that four proteins, CCDC57, MYH1, TNFAIP3 and SLC11A1, have qualifying sequence matches near a protein end and are also found in the cytoplasm, but not the nucleus. The four are therefore suggested for CR function. The remaining matches between MLARLVSNS and a human protein fail the other tests for inclusion in the CR group because the peptide is not located at a match protein end or the protein has evidence of localization in the nucleus (Table 2, lines 5-19). For instance, while the match of TTC22 with MLARLVSNS is found near the C-terminus, the protein is found in the nucleus (line 6). A similar situation is observed with PDZD7 (line 19).

**Table 2:**
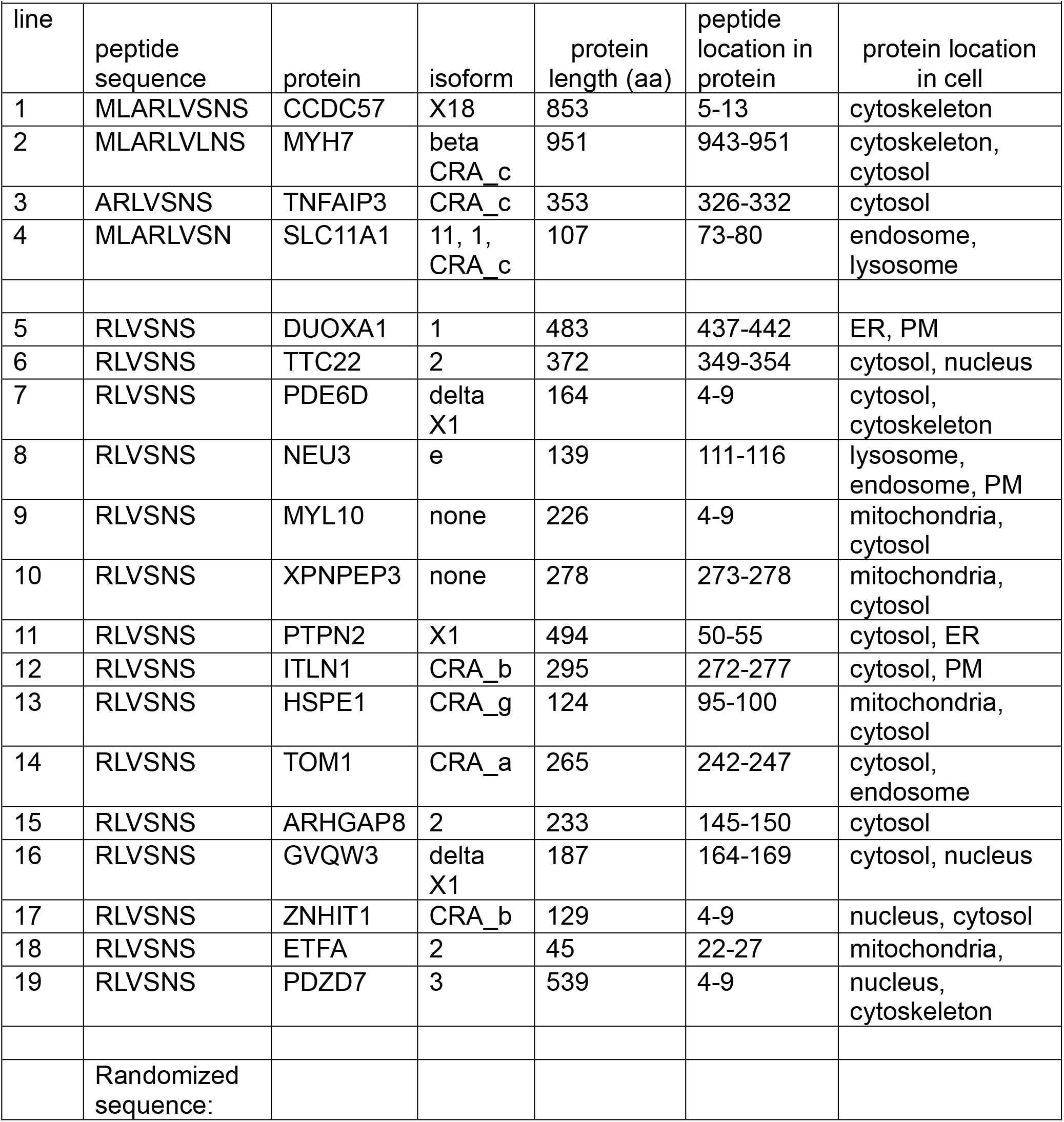
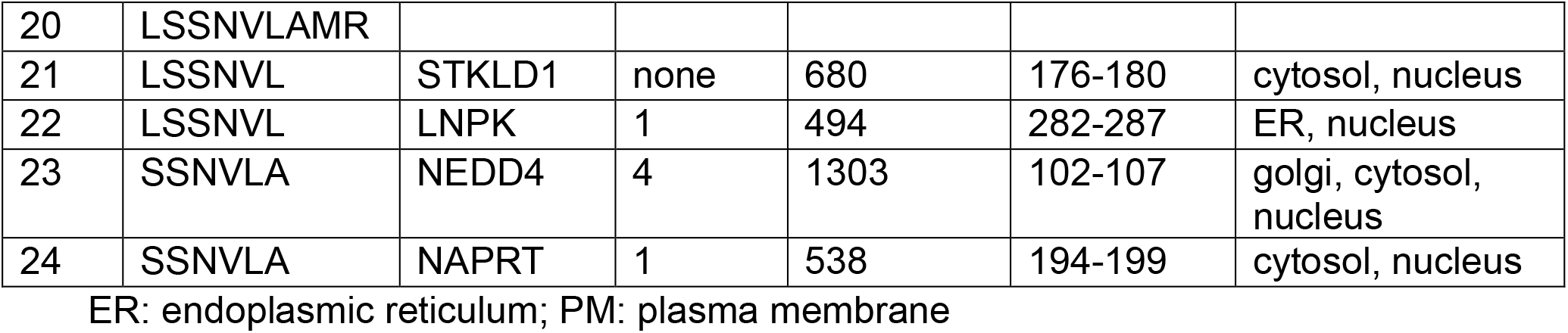
Identification of human proteins with proposed cytoplasmic retention sequence MLARLVSNS.

The BLAST results with MLARLVSNS stand out because 14 of the 19 sequence matches with human proteins involve the same 6 amino acid sequence, RLVSNS (lines 5-19). This result was not expected as three other contiguous 6 amino acid sequences are found in the test peptide (i.e. ARLVSN, LARLVL, and MLARLV) and would have a chance of being found in human genes. It is suggested that the missing sequences may have an unknown toxicity or, alternatively, that RLVSNS-containing proteins could have additional properties that extend the abundance of RLVSNS.

As in the case of the MLPRLVLNS study, a control analysis was carried out with a randomized version of MLARLVSNS using the same BLAST procedure used for the unmodified sequence (Table 2, lines 20-24). The results with randomized MLARLVSNS yielded a lower number of matches with human genes than the unmodified sequence. Four matches were found with the randomized version compared with 19 in the original one (see the protein column in Table 2). Other features of the randomized sequence, such as location in the nucleus disqualify the randomized version for inclusion in the candidate CR group.

### Protein isoform analysis

Isoforms of proteins with candidate CR sequences were examined in case they would be found to have candidate CR sequences different from the original isoforms identified. Novel candidate CR sequences were found in isoforms of five of the proteins described above to have proposed CR signals, MAP3K5, TRPC1, TTPA, CCDC57 and MYH7 (Tables 3 and 4). In each case the CR sequence found in the new isoform was observed in the same location as that in the original peptide. This situation is illustrated in the case of MAP3K5 (Table 3). Here the original test peptide sequence, MLPRLVLNS, is found in two isoforms, X8 and X9 where it is located 18 amino acids from the C-terminal end (see Table 3, lines 1 and 2). In agreement, the novel peptide DLKCLRLRG is found in isoforms 1, 2 and 6 where it is also located 18 amino acids removed from the C-terminal end. This similarity is observed even though isoforms X8/X9 and 1/2/6 differ substantially in length (i.e. ∼800aa in X8/X9 vs. 1155-1465aa in isoforms 1, 2 and 6; see Table 3 and Table 5). The same identity of isoform and proposed CR location was observed with isoforms of TTPA. Here each of the three isoforms examined has a different candidate CR peptide yet each is located at the C-terminus of the protein (Table 3, lines 9-11). The results are interpretated to indicate that the location of the CR sequence is the same in all isoforms of a particular protein while isoforms may be the same or different in the sequence of their CR peptide.

**Table 3:**
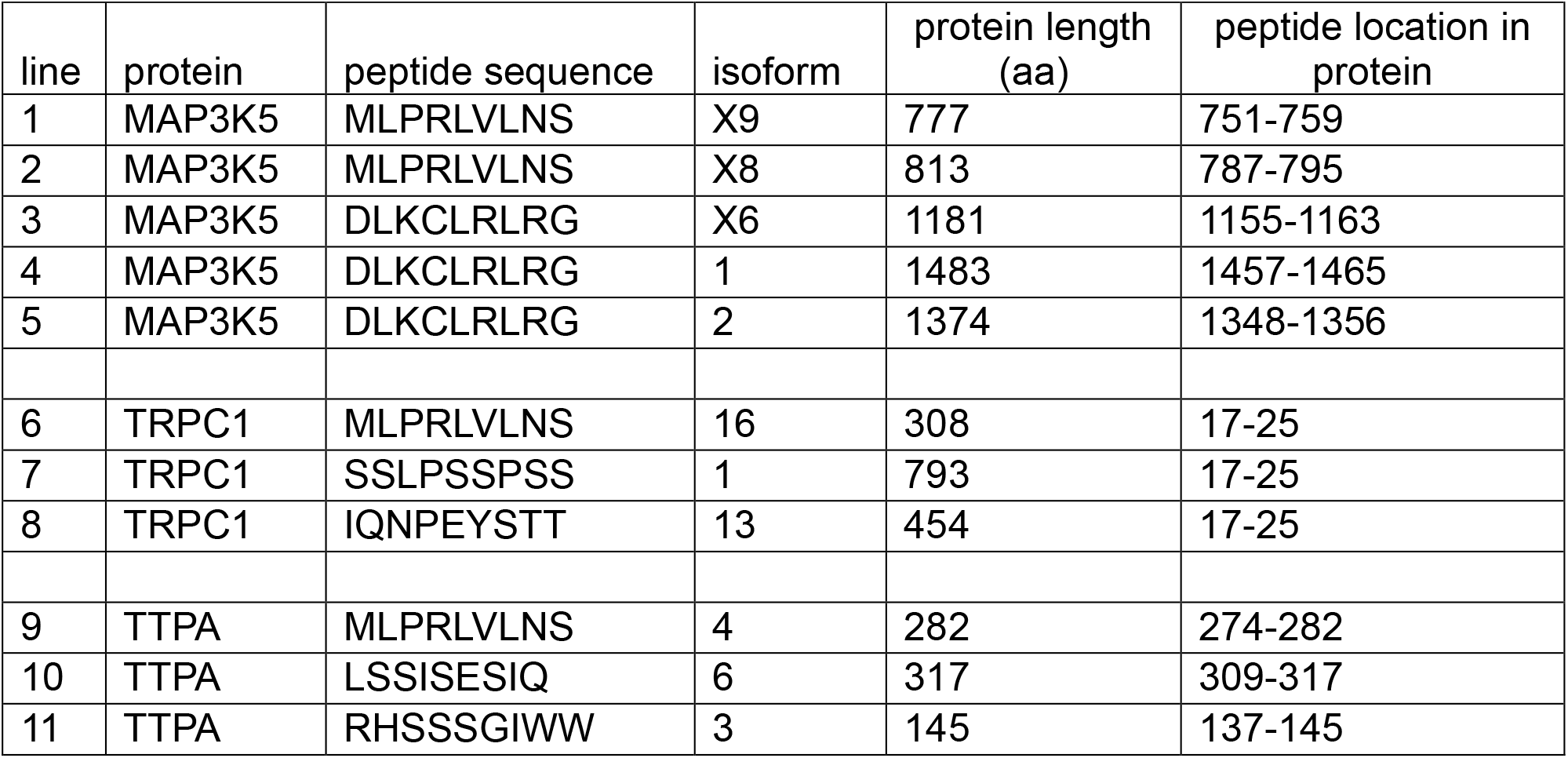
Proposed cytoplasmic retention sequences related to MLPRLVLNS in isoforms of human proteins.

**Table 4:**
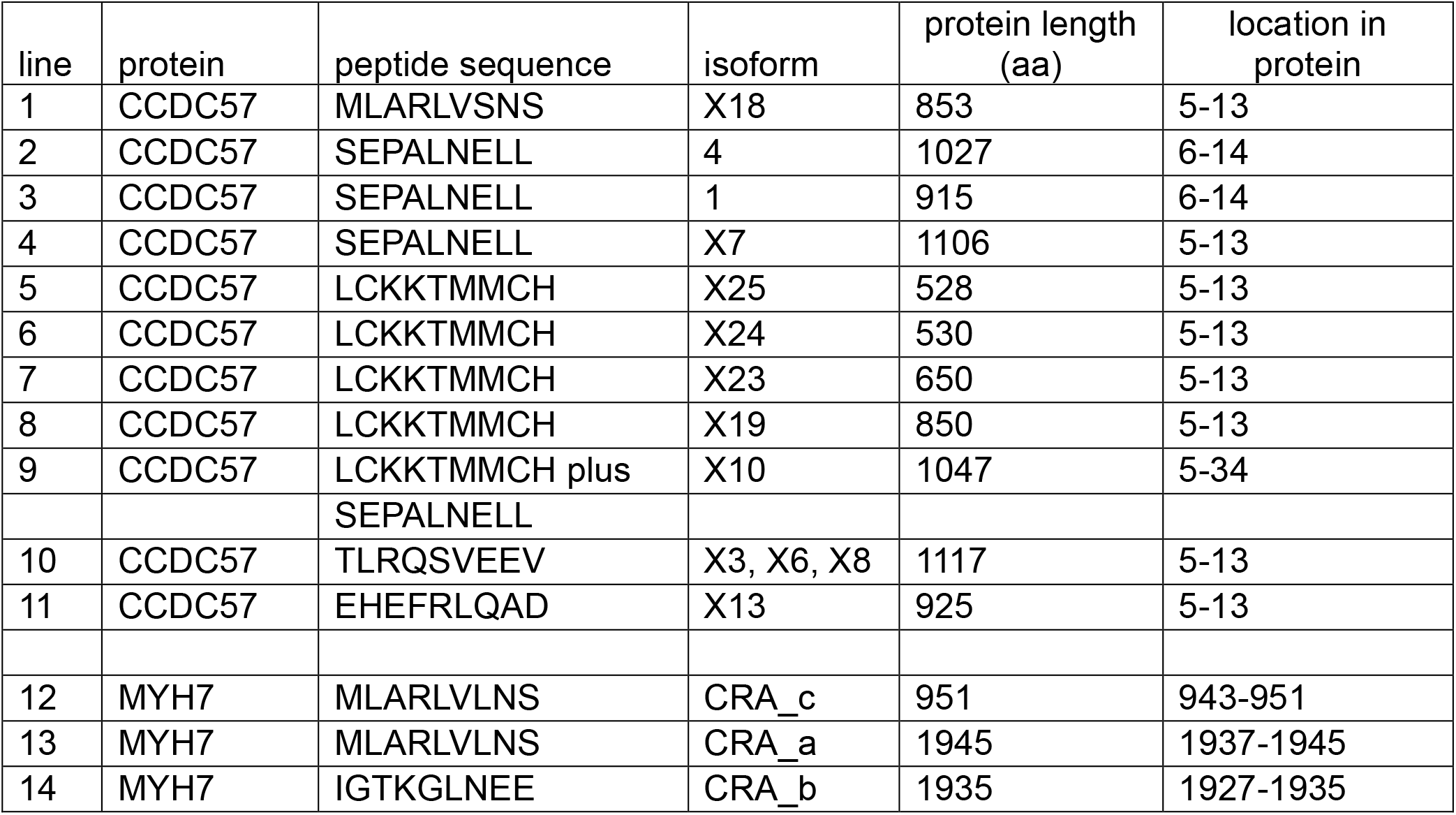
Proposed cytoplasmic retention sequences related to MLARLNSNS in isoforms of human proteins.

**Table 5:**
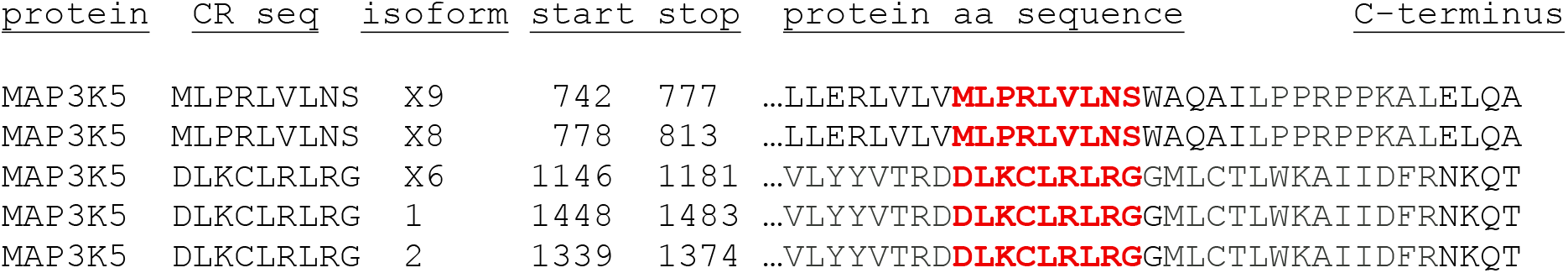
Isoform and CR sequence location in human MAP3K5.

The highest number of novel CR peptides was observed in CCDC57 (Table 4). Here a total of five distinct proposed CR sequences is distributed among 13 isoforms. In each isoform, the proposed CR sequence is located at the same position in the CCDC57 protein beginning 5-6 amino acids from the N-terminus. An unusual feature was observed in CCDC57 isoform X10 where two different, novel CR sequences are found at the N-terminal site usually occupied by only one. Here both LCKKTMMCH and SEPALNELL are found between positions 5 and 34 (line 9).

### Predicted structures of candidate CR peptides

The analysis described above yielded identification of 15 novel, candidate CR peptide sequences (Table 6). The list is noteworthy for the amino acid sequence heterogeneity observed. Heterogeneity was observed in all but five of the candidate CR sequences identified here (Table 5, lines 1, and 7-10). Apart from the five peptides, however, it is difficult to identify sequences that have obvious regions of similarity. The observed sequence heterogeneity suggests a corresponding heterogeneity in peptide structure, and this feature was examined by using AlphaFold to predict the structure of each peptide. A sample of the results is shown in Figure 1.

**Table 6:**
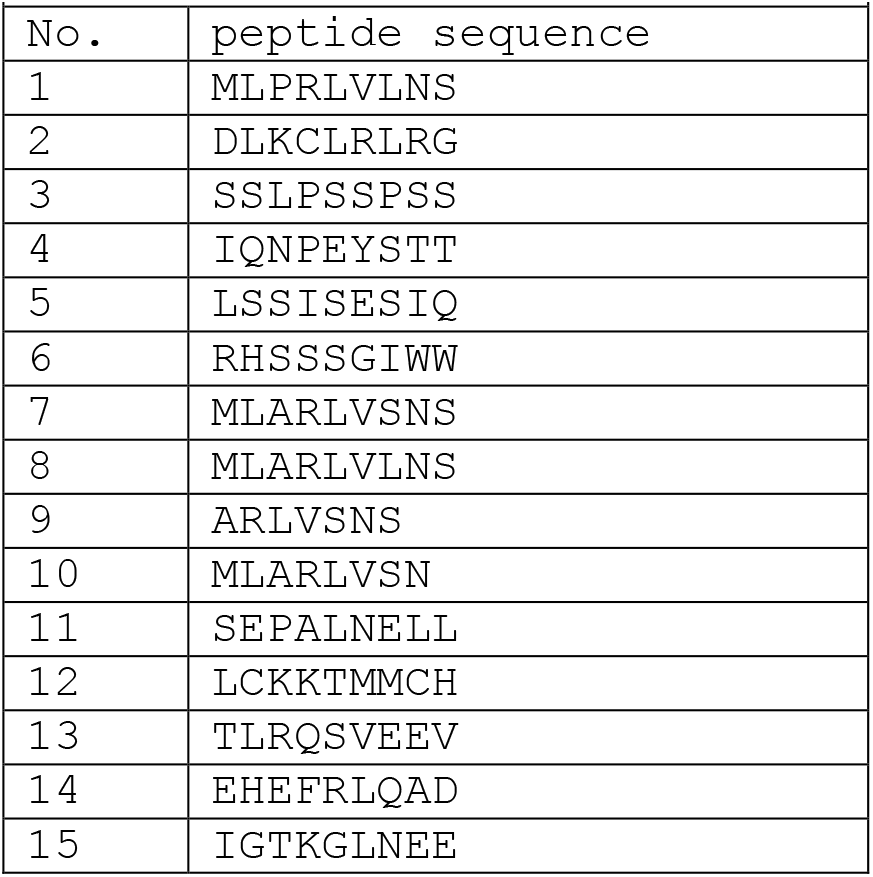
Candidate CR peptides reported here.

| Table 6: Candidate CR peptides reported here |  |
| --- | --- |
| No. | peptide sequence |
| 1 | MLPRLVLNS |
| 2 | DLKCLRLRG |
| 3 | SSLPSSPSS |
| 4 | IQNPEYSTT |
| 5 | LSSISESIQ |
| 6 | RHSSSGIWW |
| 7 | MLARLVSNS |
| 8 | MLARLVLNS |
| 9 | ARLVSNS |
| 10 | MLARLVSN |
| 11 | SEPALNELL |
| 12 | LCKKTMMCH |
| 13 | TLRQSVEEV |
| 14 | EHEFRLQAD |
| 15 | IGTKGLNEE |

**Fig. 1:**
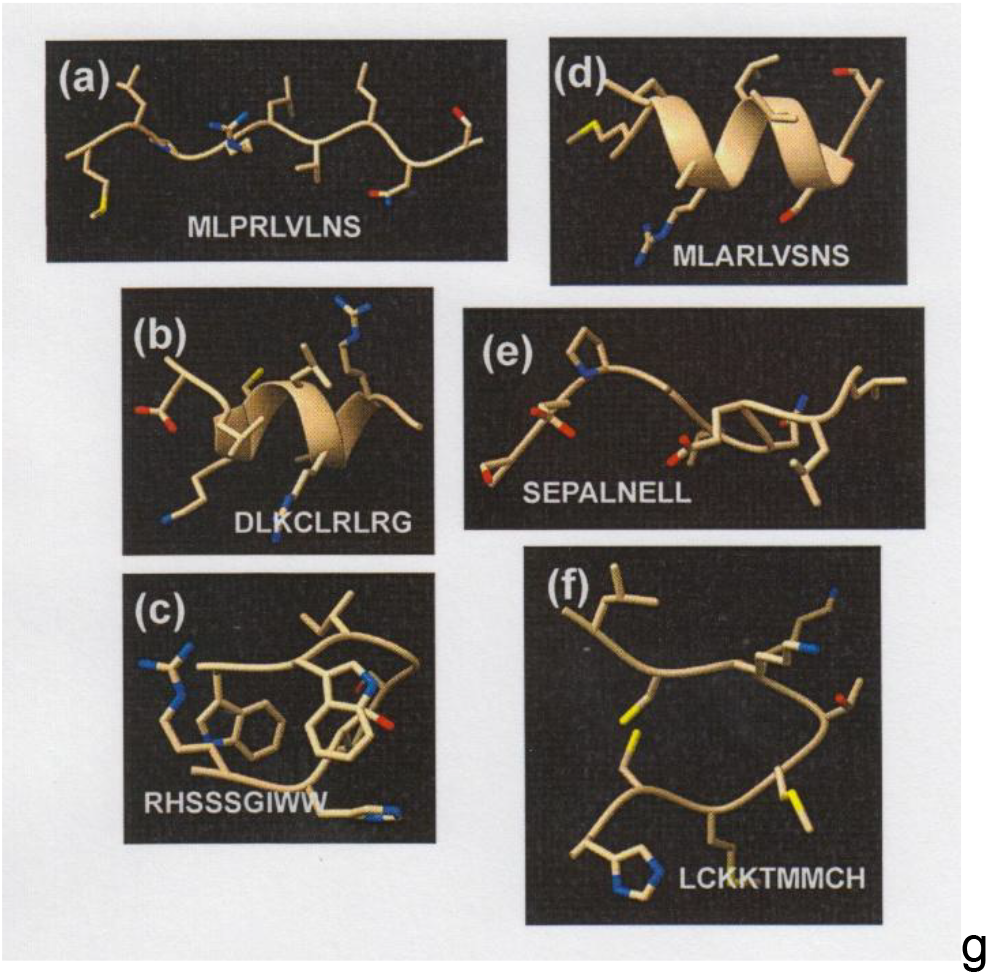
Predicted structures of selected candidate cytoplasmic retention sequences identified here. Note that the predicted structures are diverse as expected from their diverse amino acid sequences and their expected function in cytoplasmic retention.

Most of the structures could be assigned to one of three groups: (1) an extended array of amino acids as found in MLPRLVLNS and SEPALNELL; (2) a short α-helix (MLARLVSNS and DLKCLRLRG) and (3) a structure in which the peptide is folded into a U shape shown with the closed end shown at the right in RHSSSGIWW and LCKKTMMCH. In all three cases the structures are interpreted to support the view that they are highly distinct from each other. This can be seen in the α-helical pair shown in Figure 1. While DLKCLRLRG is found to be rich in positively-charged amino acids, MLARLVSNS is less so (compare panels (b) and (d) in Figure 1). DLKCLRLRG has regularly-spaced leucine residues characteristic of α-helices [9]. In the extended chain pair, SEPALNELL is found to have a hydrophobic C-terminus not found in the other extended chain structure (compare panels (a) and (e)). In the LCKKTMMCH structure two cysteine residues are near each other and potentially able to form a disulfide bond not seen in the other structures. The results of the predicted structures are interpreted to be consistent with the diverse sequences of the peptides examined and with the function of the peptides in CR of the parent protein.

## 4. Discussion

The major contribution described here is a bioinformatic method to identify protein amino acid sequences involved in retaining their parent protein in the cytoplasm. In their environment in the cell, each sequence identified is expected to be a part of a donor-acceptor pair in which both proteins spend at least a part of their lives in the cytoplasm. As there are many such cytoplasmic proteins, it is expected that there are many amino acid sequences devoted to getting proteins to the right place in the cytoplasm (e.g. the mitochondria) and keeping them in place unless movement serves another requirement of the cell. The method described here is expected to provide investigators with a rapid process to identify candidate CR sequences by focusing on their basic properties, presence of the parent protein in the cytoplasm and location of the CR sequence at an end of the protein amino acid sequence. To validate the proposed method, it has been used to identify fifteen novel candidate CR sequences in human proteins. All have the properties expected of a CR sequence including the observation that the group is quite diverse in amino acid sequence as would be expected of a small number of sequences selected from a much larger pool. It is hoped that the new method will enable further characterization of CR sequences including information about the nature of the binding sites recognized by CR sequences.

Of the observations reported here, the most consequential may be those regarding isoforms of CR containing proteins. It is observed that while different isoforms of a cytoplasmic protein may have CR sequences that are the same or different, the location of the CR sequence is the same in all isoforms. This was found to be the case even in the protein CCDC57 that has the most isoforms (13) and the most distinct CR sequences (5) of the proteins examined. Although other explanations may apply, we assume this arises by alternative splicing of the relevant protein followed by evolutionary adjustment of the isoform length so that the CR sequence can be accommodated by its binding site in the cytoplasm [10].

## AUTHOR CONTRIBUTIONS

Both authors were involved in all aspects of the research described here.

